# Structure of an infectious mammalian prion

**DOI:** 10.1101/2021.02.14.431014

**Authors:** Allison Kraus, Forrest Hoyt, Cindi L. Schwartz, Bryan Hansen, Andrew G. Hughson, Efrosini Artikis, Brent Race, Byron Caughey

## Abstract

Classical mammalian prions are assemblies of prion protein molecules that are extraordinarily transmissible, with a microgram of protein containing up to 10^8^ lethal doses of infectivity^1,2^. Unlike most other pathologic and amyloidogenic proteins, prions typically contain glycolipid anchors ^3^ and abundant asparagine‐linked glycans^4‐6^. The infectious nature, complexity, and biophysical properties of prions have complicated structural analyses and stymied any prior elucidation of 3D conformation at the polypeptide backbone level^7^. Here we have determined the structure of the core of a fully infectious, brain‐derived prion by cryo‐electron microscopy with ∼3.1 Å resolution. The purified prions are amyloid fibrils comprised of monomers assembled with parallel in‐register intermolecular beta sheets and connecting chains. Residues ∼95‐227 of each monomer provide one rung of the ordered fibril core, with the glycans and glycolipid anchor projecting from the lateral surfaces of the fibril. The fibril ends, where prion growth occurs, are formed by single monomers in an extended serpentine combination of β‐ arches, a Greek key, and loops that presumably template the refolding of incoming monomers. Our results describe an atomic model to underpin detailed molecular hypotheses of how pathologic prion proteins can propagate as infectious agents, and how such propagation and associated pathogenesis might be impeded.

## Main

Mammalian prion diseases include Creutzfeldt‐Jakob disease, bovine spongiform encephalopathy, chronic wasting disease, and scrapie^8,9^. These and many other prion diseases are transmissible, largely untreatable, and fatal. During prion infections, prions can multiply in the host by many orders of magnitude. Although it has long been apparent that prions have high β‐sheet content^1,10,11^ and propagate via templated conformational conversion of the host’s normal prion protein (PrP) isoform, PrP^C 12‐14^, the detailed 3D structures that make this happen have been elusive. PrP‐based prions are comprised primarily of misfolded multimers^15,16^, generically called PrP^Sc^ ^8^. PrP^C^ exists primarily as a heavily glycosylated, glycophosphatidylinositol (GPI)‐linked and membrane‐bound monomer^3‐6,17^ with a predominantly α‐helical C‐terminal domain and an intrinsically disordered N‐terminal domain^18^. Upon conversion to PrP^Sc^, refolding into high‐β‐sheet conformers^1,10,11^ occurs with assembly into ordered aggregates such as amyloid fibrils^19,20^ and 2D crystals^21^. Although the most infectious prion particles per unit protein were found to be ∼600 kDa, much larger amyloid fibrils are also highly infectious^16^. PrP^Sc^ multimers can induce, or seed, the conversion of PrP^C^ into PrP^Sc^ ^12,22^ in a manner that maintains prion strain‐specific PrP^Sc^ conformations^13^.

Although the characteristics of *bona fide* PrP^Sc^ fibrils have foiled any previous determination of their detailed 3D structures, low‐resolution analyses have been interpreted as indicating 2 protofilaments^19,23‐ 27^. Such studies supported a plausible, but as yet unvalidated, model in which protofilaments comprised of stacked 4‐rung β‐solenoid monomers are loosely intertwined to give a cross‐section with 2‐fold axial symmetry^25,26^. In contrast, synthetic amyloid fibrils of full‐length or N‐ and/or C‐terminally truncated bacterially‐derived recombinant PrP can adopt parallel‐in‐register intermolecular β‐sheet (PIRIBS)‐based architectures^28‐34^. While two types of recombinant PIRIBS‐based fibrils have shown detectable infectivity when large amounts were injected into the brains of rodents^35,36^, the observed attack rates and/or long incubation periods suggest that these fibrils had extremely low titers per unit protein relative to most natural prion strains. Nonetheless, these and other empirical findings have supported hypothetical PIRIBS‐based models for PrP^Sc^ in which a single PrP molecule spans the entire fibril axis to give an asymmetric cross‐section^31^.

Importantly, following recent seminal cryo‐EM‐based structure determinations for various non‐PrP protein amyloids (e.g.^37^), high‐resolution cryo‐EM structures of synthetic recombinant human PrP94‐178 (rhu94‐178)^33^ and PrP23‐231 (rhuPrP23‐231)^34^ amyloid fibrils were recently solved. Each of these fibrils have 2 protofilaments with PIRIBS‐based proteinase K (PK)‐resistant cores that are much smaller than *bona fide* infectious PrP^Sc^ fibrils and, therefore, are likely to lack infectivity^38,39^. Thus, although diverse types of evidence have constrained features of PrP fibril structures and their variants, no data from fully infectious prions have revealed the folding of monomers within PrP^Sc^ assemblies, or allowed unambiguous discrimination between widely divergent hypothetical prion models^7,9,26,31^. Here, we report a high‐resolution cryo‐EM structure of highly infectious, brain‐derived PrP^Sc^ fibrils with GPI anchors and the full complement of asparagine (N)‐linked glycans^3,6^.

### PrP^Sc^ purification, infectivity, seeding activity, and negative stain EM

We purified PK‐resistant PrP^Sc^ fibrils from brains of clinically ill hamsters infected with the 263K scrapie prion strain. Gel electrophoretic analyses of the fibril preparations indicated predominant bands at ∼20‐ 32 kDa, or oligomers thereof, that comigrated with bands reacting with PrP antibodies in immunoblots (Extended Data Fig. 1a). As expected, PK‐treatment caused a size shift consistent with cleavages within the N‐terminal domain up to ∼residue 90, leaving the remaining C‐terminal residues intact^40,41^. Densitometry of protein‐stained gels indicated that the preparations were ∼97‐98% PrP. Incubation time bioassays yielded estimated titers of 5.0 × 10^8^ to 2.0 × 10^9^ 50% lethal doses per mg protein (Methods and Extended Data Fig. 1d,e). Prion seeding assays^42^ showed that our final manipulations immediately prior to preparation of EM grids (see Methods) did not affect seeding activity (Extended Data Fig. 1b,c). Negative stain transmission EM of these preparations indicated fibrillar structures (Fig. 1a). Although the fibrils were often bundled and matted, the many isolated fibrils that we found had twists with narrow cross‐over points.

**Fig. 1.**
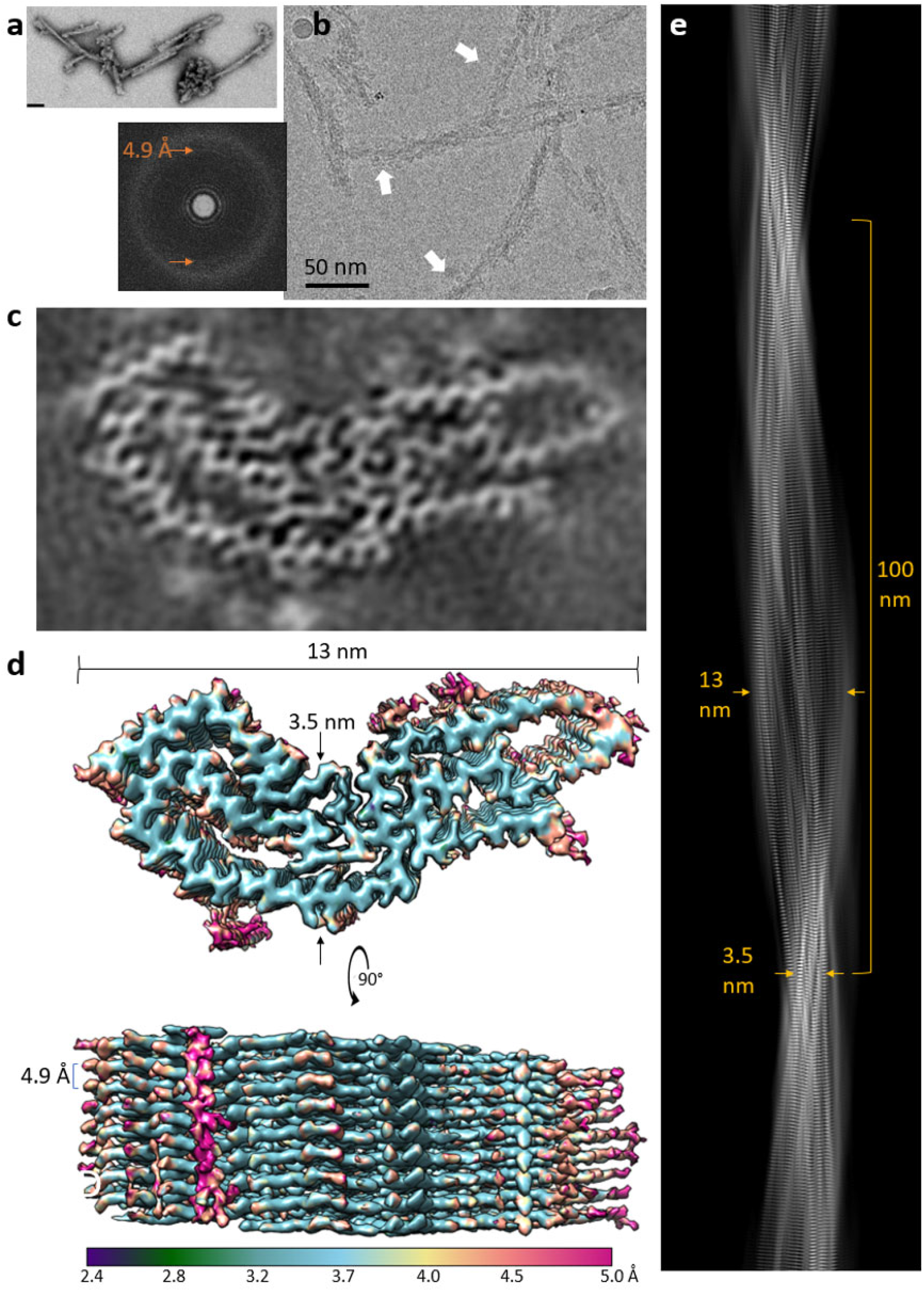
EM images and density maps of 263K prion fibrils. **a**, Negatively stained transmission EM image (bar = 50 nm). **b**, Raw 2D cryo‐EM image (cropped representation) with associated fast Fourier transform showing signals for regular 4.9 Å spacings (orange arrows). White arrows indicate globules. **c**, Projection of density map of fibril cross‐section derived from single particle cryo‐EM analysis. **d**, Surface depictions of density map with indicated dimensions and colors showing local resolutions according to bar. **e**, Projection of the fibril density map.

### Tomographic analyses

Cryo‐EM revealed that unstained 263K fibrils (Extended Data Fig. 2) had widths at the widest point of 13‐20 nm (Extended Data Table 1). Of the isolated 263K fibrils for which twist was determinable (n=401), >99% were left‐handed across 2 preparations (Extended Data Fig. 2e‐h & Supplementary Information Movies 1‐3). Approximately 45% of fibrils had visible ∼4 nm globular features aligned along one side. The globules were spaced at 7‐10 nm but could be seen along the entire length of some fibrils (Extended Data Fig. 2a‐b & Supplementary Information Movies 1‐3) and increased overall fibril widths by ∼2‐4 nm (Extended Data Table 1). Although the nature of these globules remains unclear, they might have been due in part to binding of residual detergent or other non‐PrP^Sc^ molecules. Nonetheless, the single‐sidedness of the globules implied an asymmetric fibril cross‐section. Importantly, such asymmetry differs from the symmetrical 2‐protofilament cores that have been shown for the rhuPrP23‐231 fibrils^34^ and rhuPrP94‐178 fibrils^33^, and postulated for mouse scrapie prions^25‐27^.

### Single particle analysis and 3D image reconstruction

Additional features of PrP^Sc^ fibrils were resolved with single particle acquisition and helical reconstruction. Although this process inevitably yields a small subset of the initial particle set, several factors suggest that our results represent the preponderance of PrP^Sc^ in our fibril preparations (see Supplementary Information). Imaging and data parameters are provided in Extended Data Table 2. Fast Fourier transforms of raw 2D cryo‐EM images gave weak signals indicating regular 4.9 Å spacings (Fig. 1b, offset); these signals were much stronger in transforms of particle 2D class averages (Extended Data Fig. 3d). Such spacings were also visible perpendicular to the fibril axis in images of 2D class averages. Cross‐over points and multiple axial bands of density defined the twist along the fibril axis (Extended Data Fig. 3b,d). 3D classifications using the 2D particle class averages ultimately converged on a single core morphology. In contrast to previous low resolution cryo‐EM‐based modelling of a mouse prion strain^25,26^, our 3D density map was reconstructed from the entire fibrillar cross‐section (rather than halves assumed to be equivalent) without the computational imposition of symmetry about the fibril axis.

Using helical reconstruction techniques, we determined a 3.1 Å resolution map of the fibril core (Fig. 1 & 2, Extended Data Fig. 3d & Table 2). Lateral views again indicated rungs with regular 4.9 Å spacing perpendicular to the fibril axis (Fig. 1d,e). The rungs were highly uniform, which is consistent with a PIRIBS‐based architecture in which each rung is the same, i.e. comprised of one PrP molecule, but inconsistent with β‐solenoid models in which adjacent rungs are formed by different PrP segments. Accordingly, we built a 3D structural model by threading the PrP polypeptide backbone through densities in the fibril cross‐section (Fig. 2b).

**Fig. 2.**
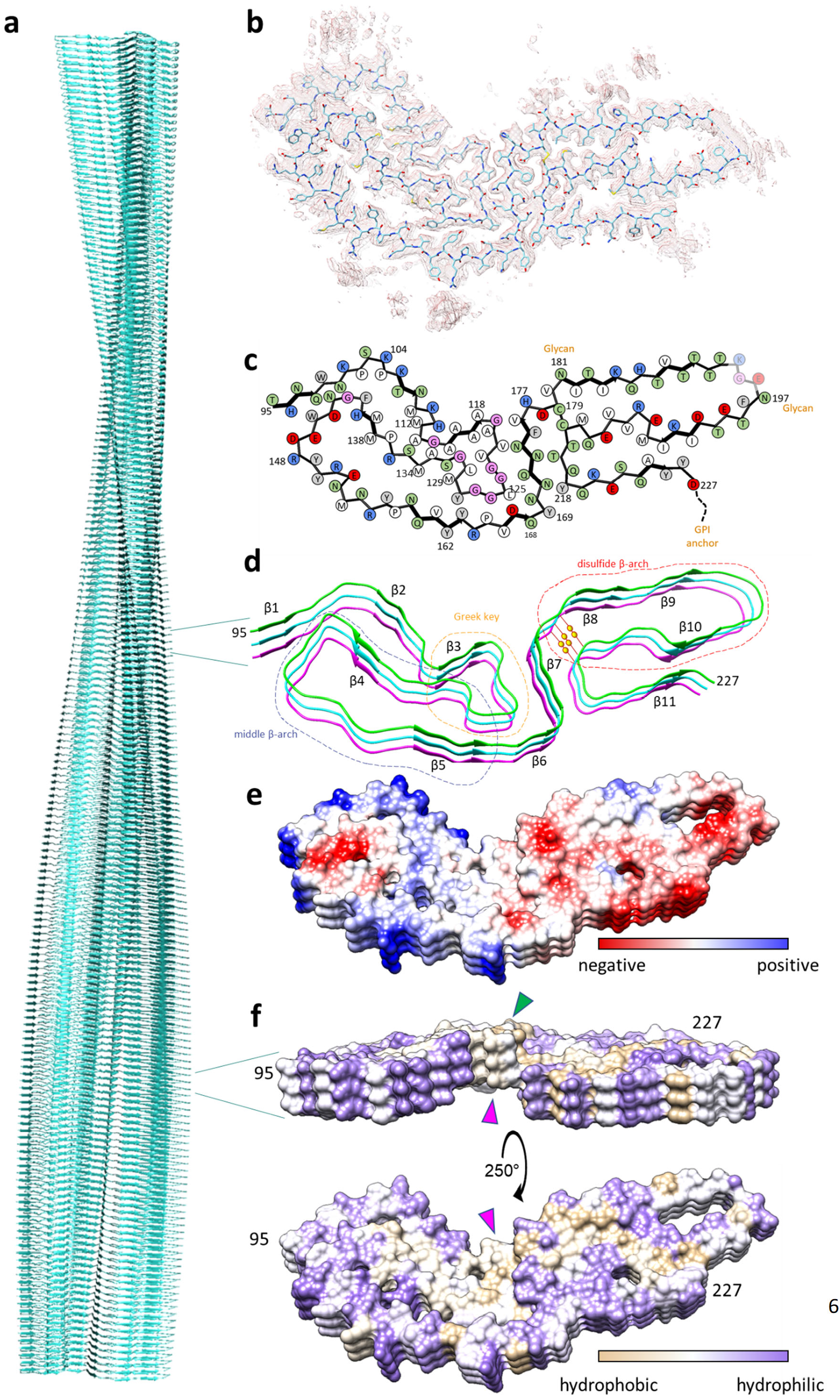
263K prion model based on cryo‐EM density map. **a**, Extended fibril model as a ribbon diagram. **b**, PrP residues 95‐227 threaded through a cross‐sectional density map (mesh). **c**, Schematic depiction of fibril core showing side chain orientations relative to the polypeptide backbone. Residues assigned to β‐sheets in Chimera are marked by thicker backbones with arrowheads. Side chains of residues 194‐196 (faded) were poorly resolved. **d**, β‐sheets in a stacked trimeric segment of the fibril. Structural elements as labelled and disulfide bond indicated by pair of yellow spheres. **e**, Coulombic charge representation. **f**, Kyte‐Doolittle hydrophobicity surface of fibril ends (templates) showing the protruding (green arrowhead) and receding (magenta arrowheads) hydrophobic Greek key motif at opposite ends.

### Cross‐sectional asymmetry

In this structure, the cross‐section is indeed asymmetric about the fibril axis (Fig. 1 & 2b‐f). The well‐resolved fibril core, which primarily represents residues 95‐227, was ∼13 nm in longest dimension and ∼3.5 nm at its narrowest (Fig. 1e). We infer from the known attachment sites of the glycans (N181 and N197) and GPI anchor (S231) that these moieties project into the solvent. However, these structures were largely unresolved in the averaged image reconstruction, presumably because of their heterogeneity in sequence^6^ and conformation^43^. Nonetheless, the first unit of N‐linked glycans, N‐acetylglucosamine, is consistent with the densities that are adjacent to N181 and N197 (Extended Data Figs. 4 & 5). Despite published doubts that bulky glycans could fit on each rung of a PIRIBS‐based core of an amyloid^7^, recent *in silico* studies have shown that such glycans can be accommodated^43,44^. We note that the GPI anchor and N197‐linked glycans are aligned asymmetrically along one edge of the fibril where they might anchor and/or contribute to the globules observed in single particle imaging and tomography (Fig. 1b & Extended Data Fig. 2). Another key asymmetric feature of the core is the preponderance of cationic residues in the N‐terminal half and anionic residues in the C‐terminal half (Fig. 2e). Surface hydrophobicity maps show hydrophobic zones in both halves, as well as a hydrophobic bridge across the interface between the two halves (Fig. 2f). On the N terminal side, a tightly packed dry zipper seals residues 96‐100 to 142‐145 (Fig. 3a). Other major features of the 263K structure include the following:

**Fig. 3.**
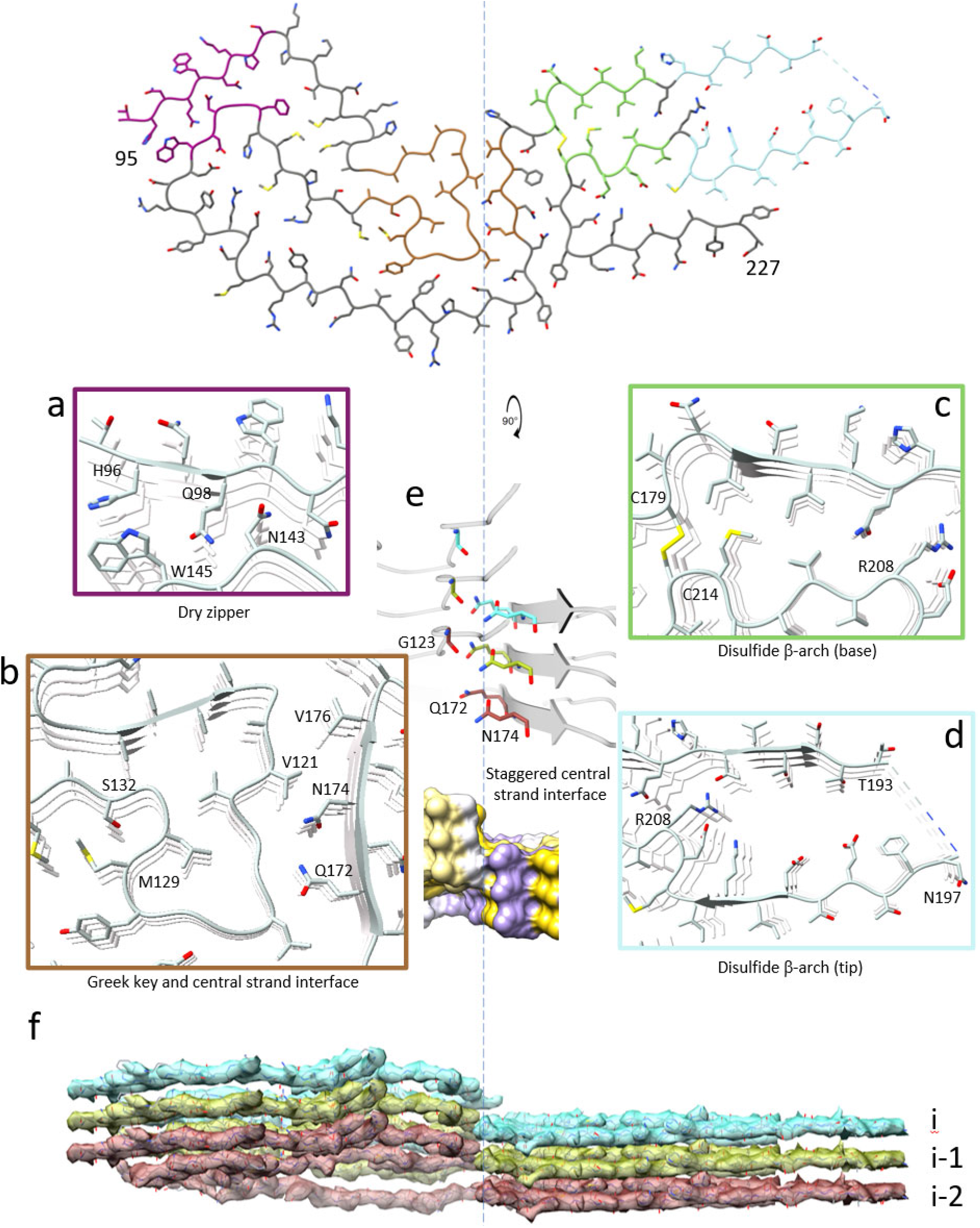
Features of the 263K prion core. **a**, Dry zipper (purple box) formed by tight interdigitation of sidechains of residues H96, Q98 and N100 with those of W145 and N143. **b**, Tip of Greek key motif (brown box) with hydrophobic packing of sidechains of A117, A120, V122 and L130 in the center and between V121 across a staggered interface (see below) to N174 and V176. **c**, Tight packing of hydrophobic sidechains at the base of the disulfide β‐arch (green box). **d**, Wide and presumably hydrated gap between the flanks of disulfide β‐arch near the tip (blue box). Dashed lines are shown in place of poorly resolved residues 194‐196. **e**, Staggered interface between Greek key residues and central strand of residues 170‐176 is shown with a ribbon diagram of interfacing residues and a hydrophobicity surface. **f**, Model (stick) and cryo‐EM map densities (transparent surface) with monomers differentiated by color.

### Hydrophobic “Greek key” motif

A particularly hydrophobic stretch of residues ∼112‐134 is organized in an architecture reminiscent of the “Greek key” topology in α‐synuclein fibrils^45^. These residues have been shown to form species‐dependent PIRIBS‐based β‐cores in rPrP23‐145 fibrils^32,46^ or a much more elongated β‐arch in rhuPrP94‐ 178 fibrils^33^. However, these previously observed topologies differ markedly from the 263K Greek key (Figs. 2b‐d & 3b and Extended Data Fig. 6a).

### Middle (125‐168) β‐arch

Residues ∼125‐168 form a major hairpin or, more conventionally, β‐arch^34^ (Figs. 2b‐d) that is related to what we had termed the “127‐161 hairpin” in earlier *in silico* modelling^31^. Our initial prediction of a β‐ arch in this region was based on an observation that the introduction of an artificial disulfide bond into the corresponding residues of murine PrP, which covalently linked the short β1 and β2 strands of PrP^C^, was compatible with conversion to PrP^Sc^ within scrapie‐infected cells^47^. However, in the 263K structure, this middle β‐arch is flipped across one or two cross‐sectional axes relative to the orientations depicted in our previous hypothetical models^31^.

### Disulfide β‐arch

263K fibrils also have a disulfide‐linked β‐arch between C179 and C214 (Figs. 2b‐d & 3c,d). This β‐arch is reminiscent of motifs that have been predicted for PIRIBS‐based prions^28,29,31^, or observed in rhuPrP23‐ 231 fibrils^34^. However, in contrast to recombinant PrP fibrils, the disulfide β‐arch in 263K prions contains two N‐linked glycans. Also, the flanks of this arch are straighter than the β‐arch in rhuPrP23‐231 fibrils^34^, except for a notable kink at M206‐R208, and, the presumably hydrated gap between flanks near the tip of the arch is larger (Fig 3d, Extended Data Fig. 6b). The extreme C‐terminal residues fold back against the flank of the disulfide loop at Y218, whereas in rhuPrP23‐231 fibrils, this fold occurs at R220. Perhaps this conformational difference is influenced by sequence differences and the GPI anchor at the C‐terminus of the 263K monomers, which is absent in rhuPrP23‐231 fibrils.

### Staggered interface between N‐ and C‐terminal domains

Lateral views of the 263K fibril show that residues of each monomer are not entirely coplanar across the fibril cross‐section and ends (Fig. 2f). At the interface between the Greek key domain and more C‐terminal residues, residues ∼119‐135 of one monomer are most closely opposed to residues 158‐177 of the i‐1 and i‐2 monomers (Fig. 3b,e,f). This results in uneven fibril ends with the hydrophobic Greek key motif protruding at one end and receding at the other (Fig. 2f).

### Further comparisons to historical data

The 263K cryo‐EM structure is consistent with a variety of other known characteristics of 263K prions or other mammalian prions that we have not already discussed above. These features include secondary structure composition, exposure (or lack thereof) of antibody epitopes, conditional exposures of minor proteolytic cleavage sites, exchange rates of PrP backbone amide protons, and the lack of clear x‐ray diffraction and cryo‐EM signals for 10 Å inter‐sheet spacings that are typical of other amyloid fibrils (see Supplementary Information and Extended Data Fig. 7).

### Conclusions and perspectives

From the cryo‐EM‐based density map of this highly infectious mammalian prion, the serpentine threading of the polypeptide backbone of residues 95‐227 and orientations of almost all of the sidechains relative to the backbone were clear. Further atomistic details of sidechain conformations were approximated by molecular modeling and best fit with the density map. The span of 134 residues packed into the ordered 263K core is more than twice as large than in the protofilament cores of previously studied recombinant PrP fibrils^32‐34^. The asymmetric 263K PIRIBS‐based core architecture is starkly different than has been proposed for murine prion fibrils based on lower‐resolution cryo‐EM and X‐ray fiber diffraction studies^25,26^, with no evidence of either β‐solenoids or independent, equivalent protofilaments. The middle (125‐168) and disulfide β‐arches were predicted in previous modelling of PrP amyloids^28,29,31^, but the relative orientation of these arches in the actual 263K prion structure is flipped relative to our earlier speculative models. The 263K disulfide β‐arch, which is heavily glycosylated, is markedly different from the unglycosylated disulfide β‐arch of rhuPrP23‐231 fibrils^34^ (Extended Data Fig. 6). Moreover, the attachment of glycans at N197 residues would block the head‐to‐head interaction observed between the 2 protofilaments in the rhuPrP23‐231 fibrils. The Greek key motif of hydrophobic residues 113‐134 is clearly distinct from those of rhuPrP23‐145^32^ or rhu94‐178^33^ (Extended Data Fig. 6). Neither this Greek key motif, the middle β‐arch, nor any other ordered folding in the N‐terminal half of the 263K prion core were resolved previously in the presumably non‐infectious rhuPrP23‐231 fibrils^34^. Thus, the additional folding in the N‐terminal half of the 263K prion core may be important for infectivity.

The C‐terminal GPI anchors twist along one side of the fibril, where they would likely bind to membranes *in vivo* (Fig. 4). This twisted tethering might often wrap the fibrils in membranes, impeding formation of amyloid plaques, obscuring fibril visualization in tissue sections, and causing membrane distortions that are pathognomonic for prion disease such as spiral membrane invaginations^17,48^. In this topology, the fibril axis must run parallel to the membrane surface, with access to much of the fibril surface blocked by either the N‐linked glycans or the membrane. The seeding surfaces at the fibril ends would then be held perpendicular to membrane surface. As postulated previously for PIRIBS‐based fibril architectures in general^31,49^, these surfaces can be envisioned to provide a precise template for the refolding of incoming monomers for the conformationally faithful propagation of PrP^Sc 13^.

**Fig. 4.**
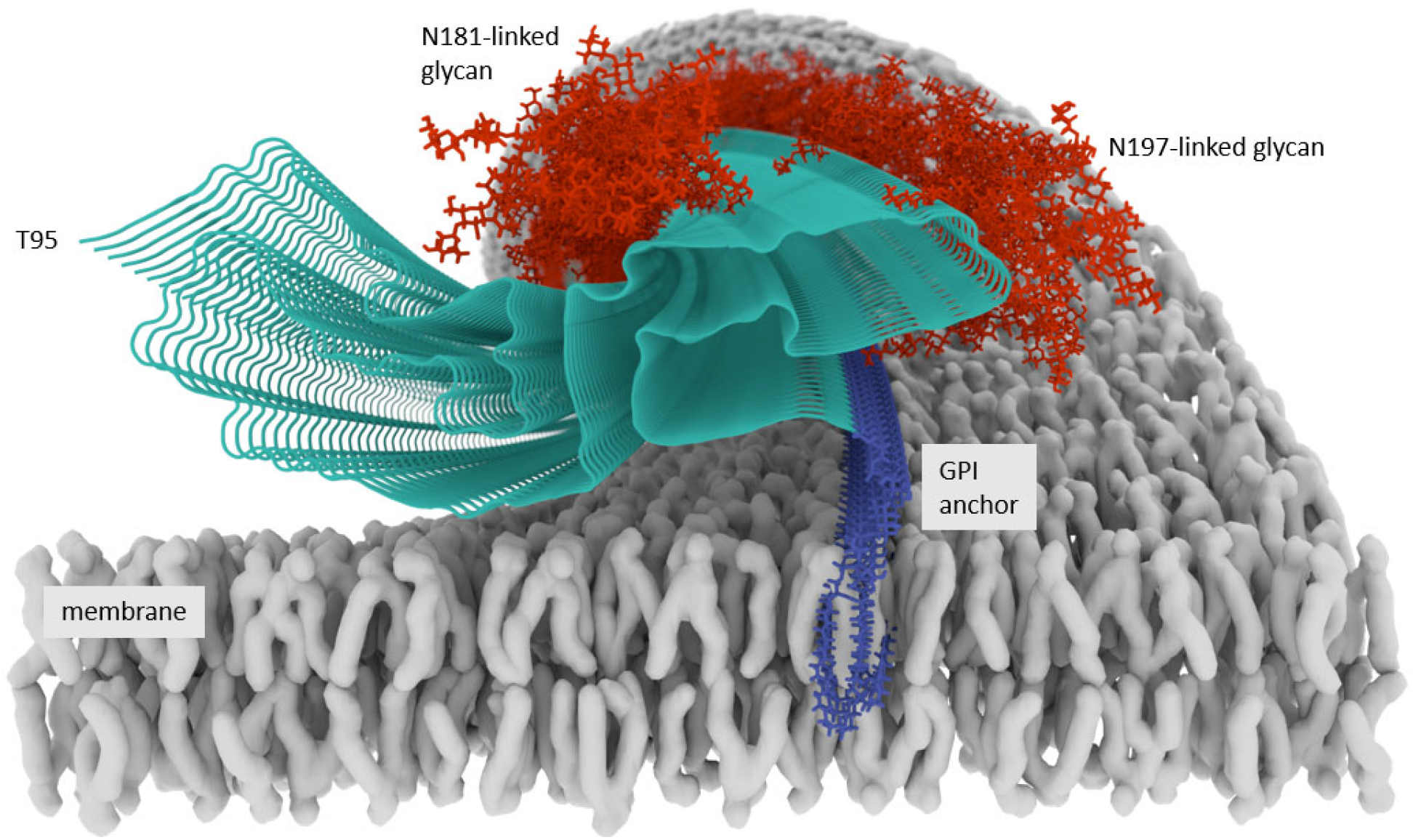
Membrane‐bound 263K prion model of ordered fibril core (turquoise) with hypothetical illustrations of glycans (red) and GPI anchors (blue) that were unresolved in the cryo‐EM images. For simplicity, single representative structures for the N‐linked glycans and GPI anchors are depicted here; however, the actual glycan and GPI anchor structures and conformations will be more varied^6^. The curl of the membrane (gray) is shown hypothetically to follow the axial twist of the GPI anchors.

Comparison of the structures of PrP^C 50^ and 263K fibrils indicates that key events in this conformational conversion are the peeling of PrP^C^’s β1‐helix 1‐β2 loop away from helices 2 and 3 and the unspiraling of each of the helices to form the extended strands of the middle and disulfide β‐arches in PrP^Sc^ (Extended Data Fig. 8). This conversion also requires disruption of the β1‐β2 sheet of PrP^C^, with the associated intramolecular hydrogen bonds becoming intermolecular in PrP^Sc^. Further studies will be needed to determine the order of these and other conformational changes, and the extent to which permutations in PIRIBS‐based architectures, and even completely different architectures such as β‐solenoids, might populate the range of infectious PrP‐based prion strains.

## Methods

### Animal studies

Animals were housed at RML in an AAALAC‐accredited facility. Experiments were in accordance with the NIH RML Animal Care and Use Committee approved protocols (2018‐011 and 2016‐039). Syrian golden hamsters were purchased from Envigo. Tg7 mice^51^ for bioassays were obtained from colonies maintained at RML. To generate 263K prions for the current study, hamsters were anesthetized and intracerebrally inoculated with 30 µl of 1% 263K scrapie brain homogenate stock containing 3.8 × 10^6^ 50% lethal doses (LD_50_). Animals were euthanized from 72‐78 days post inoculation when consistent, advanced clinical signs of prion disease were observed. Brains were collected from these animals as described previously^52^.

### 263K PrP^Sc^ fibril purification

PK‐resistant PrP^Sc^ was purified from the hamster brain (n=20‐30 per preparation) as described previously^52^. Two independent preparations (preps 1 & 2, 10 y old and fresh, respectively) were analyzed with nearly indistinguishable results in all stages of this biochemical characterization and imaging through tomography, single particle analysis, and 3D image reconstructions at resolutions down to ∼4.2 Å. However, the animal infectivity bioassay was performed only with prep 1 and, because of insufficient quantity of prep 1, the final push to collect the 12,828 movies used to improve resolution into the 3 Å range was performed with prep 2. SDS‐PAGE and immunoblotting with a polyclonal rabbit antiserum against PrP C‐terminal residues (R20)^53^ were performed essentially as described previously^54^. Immediately prior to grid preparation, fibril preparations were vortexed and allowed to sit for 10 min to gravity pellet highly bundled fibrils. Aliquots were taken from the supernatant and diluted in 20 mM Tris pH 7.4, 100 mM NaCl containing 0.02% amphipol 8‐35 and sonicated.

### Bioassays of purified 263K prions

To assess the infectivity of purified 263K fibrils used for cryo‐EM grid preparations, fibrils from prep 1 were diluted in inoculation buffer (2% fetal bovine serum in phosphate buffered balanced saline). Serial 100‐fold dilutions containing 1 µg, 10 ng, and 100 pg PK‐resistant PrP^Sc^ were intracerebrally inoculated as 30 µL volumes into Tg7 mice (n=7‐8 per group). Infectivity titers were estimated by comparing the average incubation period at each dose to a standard curve of inoculated titer vs incubation period that was generated from previous end‐point dilution assays of 263K hamster scrapie brain homogenate in Tg7 mice. Immunohistological analyses of brain tissues were performed as described^38^.

### Prion seeding assay (RT‐QuIC) analysis

Endpoint dilution analysis of the fibril preparations was conducted using RT‐QuIC with recombinant hamster PrP90‐231 substrate as described^42^.

### Negative stain electron microscopy

Ultrathin carbon on lacey carbon support film grids (400 mesh, Ted Pella, Redding, CA) were briefly glow‐discharged and floated on droplets of fibril solutions. Grids were briefly washed by sequential immersion in milliQ water before being negatively stained with Nano‐W (2% methylamine tungstate) stain (Nanoprobes, Yaphank, NY) and wicked dry. Grids were imaged at 80 kV with a Hitachi HT‐7800 transmission electron microscope and an XR‐81 camera (Advanced Microscopy Techniques, Woburn, MA).

### Cryo‐EM

C‐Flat 1.2/1.3 300 mesh copper grids (Protochips, Morrisville, NC) were glow‐discharged with a 50:50 oxygen/hydrogen mixture in a Solarus 950 (Gatan, Pleasanton CA) for 10 s. Grids were mounted in the tweezers for an EM GP2 plunge freezer (Leica, Buffalo Grove, IL) and a 3 µl droplet of 0.05% amphipol A8‐35 in phosphate buffered saline was added to the carbon surface and hand blotted, leaving behind a very thin film. The tweezers were then raised into the chamber of the plunge freezer, which was set to 22°C and 90% humidity. 3 µl of recently sonicated sample was added to the carbon side of the grid and allowed to sit for 60 s. The sample was blotted for ∼4 s followed by a subsequent 3 s drain time and then plunged into liquid ethane kept at ‐180°C. Grids were mounted in AutoGrid assemblies and then loaded into a Titan Krios G3i (Thermo Fisher Scientific, Waltham, MA) with a K3 camera and BioQuantum GIF (Gatan, Pleasanton, CA). Images were acquired at 0.55 Å/pixel at Super Resolution mode, 60 e‐/Å2, and 60 total frames. A total of 12,828 movies were collected using SerialEM^55^.

### Cryo‐electron tomography

Grids were prepared exactly as those prepared for Cryo‐EM, except 1 µl of 5 nm Protein A gold (CMC, Utrecht, Netherlands) was added with 2 µl of sample on the grid before the 60 s wait time. In some cases, sample and gold fiducials were added to ultrathin carbon on lacey carbon support film (200 mesh, Ted Pella, Redding, CA) grids that were not treated with 0.05% amphipol, blotted by hand, and plunged into liquid ethane. Tilt‐series were acquired bidirectional from zero every 3° between ±60° for a total dose of ∼80 e^‐^/Å^2^ at a pixel size of 2.87 Å using Tomography 4 software with a Falcon III camera operating in linear mode on a Krios G1 transmission electron microscope (Thermo Fisher Scientific, Waltham, MA) operating at 300 kV. Tomograms were generated with weighted back projection using IMOD ^56^.

### Image processing

Motion correction of raw movie frames and helical reconstruction were performed with RELION 3.1^57^. CTF estimation was performed using CTFIND4.1^58^. Fibrils were picked manually (Extended Data Fig. 3a), and segments were extracted with an inter‐box distance of 14.7 Å with a 370 Å box size. A second set of particles was extracted with a box size of 1280 pixels and was downscaled to a box size of 256 pixels. Reference‐free 2D class averaging (Extended Data Fig. 3b) was performed on both particles sets using a regularization parameter of T = 2, in‐plane angular sampling rate of 2°, a tube diameter of 200 Å, and the translational offset limited to 4.9 Å. 2D classes, from the large box size, were used to estimate the cross‐over distance of the fibril. An initial 3D model was then generated, from the small box size, from multiple 2D class averages^57^. This initial model was used for 3D auto refinement with C1 symmetry, initial resolution limit of 40 Å, initial angular sampling of 3.7°, offset search range of 8 pixels, initial helical twist of ‐0.7°, initial helical rise of 4.8 Å, and using 20% of the segment central Z length. The output from auto refinement was used for 3D classification without allowing for image alignment to remove poorly aligned segments from auto refinement. Classes were selected for further refinement based on similarity of features in their cross‐section (excluding visually low resolution and poorly aligned classes), estimated resolution, overall accuracy of rotation and translation, and Fourier completeness (Extended Data Fig. 3c). Auto‐refinement was then performed while optimizing the helical twist and rise. Auto‐refinement with refinement of twist and rise yielded a final map with a twist of ‐0.847° and rise of 4.874 Å. Iterative cycles of CTF refinement, Bayesian polishing, and auto refinement were used until resolution estimates stabilized. Post processing in RELION was performed with a soft‐edged mask representing 10% of the central Z length of the fibril. Sharpening was applied with a B‐factor of ‐31 Å^2^. Resolution estimates were obtained between independent refined half‐maps at 0.143 FSC (Extended Data Fig. 3d).

### Model building

De novo building of the atomic model was carried out using Coot^59^. Assuming inclusion of residues ∼90‐ 231 into the polypeptide backbone, residue placement was initially guided by the Cys179‐Cys214 disulfide bond and nearby bulky residues of 172‐175 and 162‐164. The remaining N and C‐terminal amino acids were manually added to the model followed by targeted real‐space refinement in Coot. Individual subunits were translated along the axis of the fibril to generate a stack of three consecutive subunits to maintain interactions between adjacent subunits. The translated subunits were rigid‐body fit in Coot. Iterative real‐space refinement and validation with Coot and Phenix^60,61^ were carried out in the absence of any imposed model restraints with further local refinement in Fourier space using RefMac5^62^. Model validation was carried out with CaBLAM^63^, MolProbity^64^, and EMringer^65^, and any outliers/clashes identified and corrected with subsequent iterative refinements/validation as described above. Additional details are as listed in Extended Data Table 2.

## Supporting information

Movie 1 fibrils with globules

Movie 2 fibrils with no globules

Movie 3 fibrils whole tomogram

## Acknowledgements

We are very grateful to Prof. Sjors Scheres for his early advice and encouragement. We thank Dr. Kunpeng Li at the CWRU cryo‐EM facility for assistance in collecting cryo‐EM data. We thank Ms. Elizabeth Fisher for helpful suggestions and oversight of the EM facility; Dr. Dave Dorward for help with preliminary negative stain TEM; Drs. Suzette Priola, Bradley Groveman, Amitava Roy and Ankit Srivastava for their helpful in‐house review of this manuscript; Katie Williams for help with bioassays; Dr. Jacqueline Leung for assistance in analyzing related tomography data; Nancy Kurtz and Lori Lubke for neuropathological slide processing; and Austin Athman and Anita Mora for graphics arts assistance. This work was supported by the Intramural Research Program of the NIAID; Mary Hilderman Smith, Zoë Smith Jaye, and Jenny Smith Unruh in memory of Jeffrey Smith; and the Britton Fund, Case Research Institute, and CWRU School of Medicine. This work utilized the computational resources of the NIH HPC Biowulf cluster (http://hpc.nih.gov).

## Author information

### Contributions

AK and BC supervised the project. All authors designed experiments. AH purified the PrP^Sc^. AH and AK provided *in vitro* analyzes of PrP^Sc^ preparations. BR bioassayed PrP^Sc^ and performed neuropathological analyses. CS and AK prepared EM grids and collected EM data. CS analyzed the tomographic data. FH, CS, BH, and AK processed the cryo‐EM images. FH, AK developed map parameters and FH performed class averaging and refined EM map densities in Relion. FH, BC, AK built *de novo* atomic models of PrP^Sc^, FH and AK performed iterative refinements and map/model validation. EA performed validations for model stability and built and evaluated glycosylated and GPI‐anchored models. All authors helped to prepare the manuscript.

**Supplementary Information** is available for this paper.

## EXTENDED DATA

**Extended Data Fig. 1.**
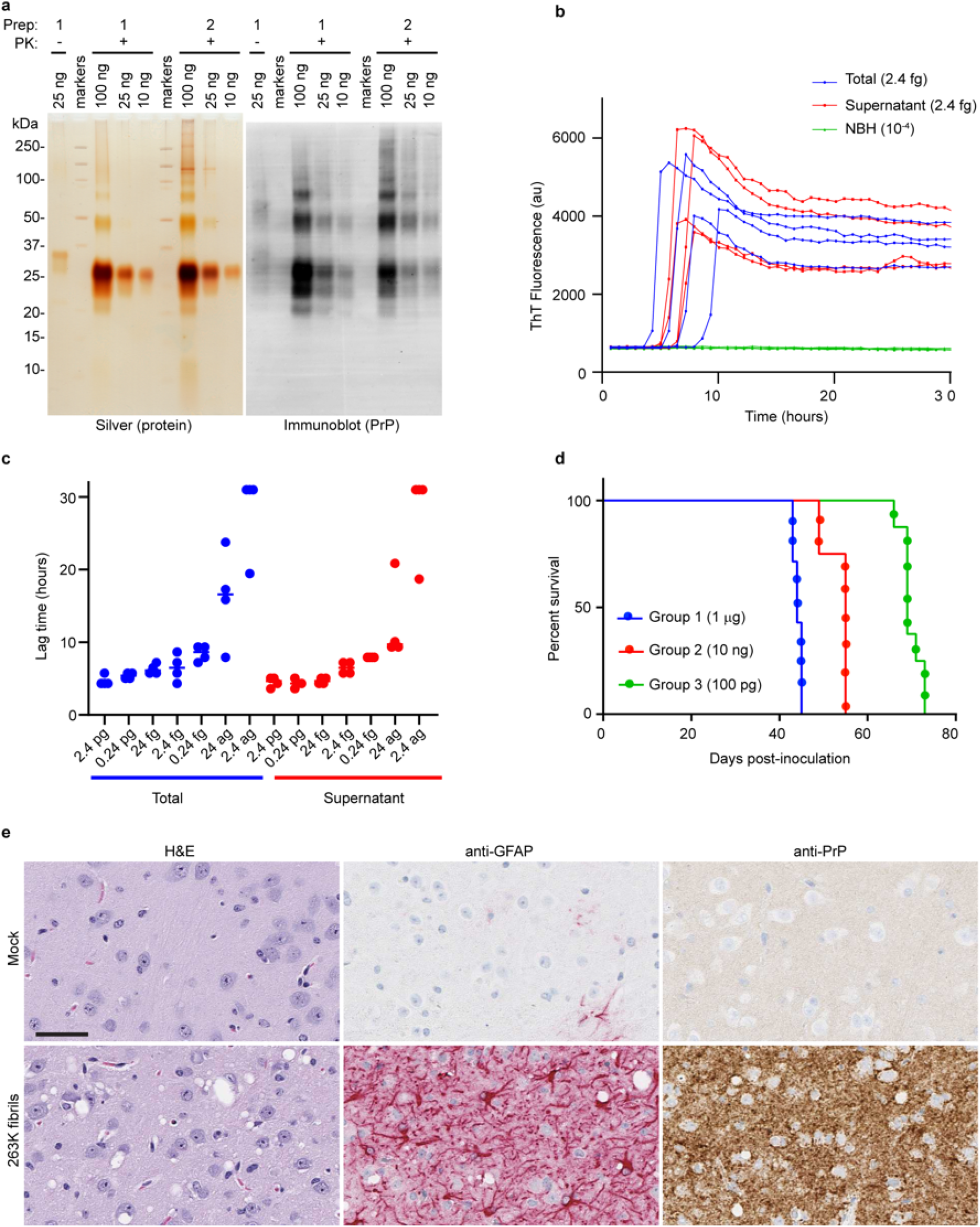
263K PrP^Sc^ preparations are protease‐resistant, seed‐competent, and highly infectious. **a**, Gel analysis of two independent 263K purifications. Purified 263K prions from two preparations were subjected to PK digestion (PK+) and used for gel analysis and subsequent silver staining or PrP immunoblotting as indicated. Untreated 263K (PK‐) is shown in the first lane. **b**, Traces from quadruplicate ThT readings over time for RT‐QuIC reactions seeded with 2.4 fg of either the initial purification of prep 1 (Total), or the same subjected to subsequent manipulations to improve fibril distribution (Supernatant). Negative control reactions seeded with normal brain homogenate (NBH) at 10^‐4^ tissue w/v dilution. **c**, Endpoint dilution RT‐QuIC comparison of lag times of individual reaction wells (n=4) seeded with the Total or Supernatant fractions of 263K prep 1. Data points above 30 h represent reactions that failed to cross the positivity threshold within the total 30‐h reaction time. Bar indicate means. **d**, Kaplan‐Meier survival curves after inoculation of the designated amount of 263K fibril prep 1 in transgenic mice (n = 7‐8 per group) that overexpress hamster PrP^C^. Survival time was determined by the need to euthanize the mice according to clinical criteria for definite scrapie as referenced in Methods. Dots represent individual animals euthanized at the designated time. **e**, Brain tissue from animals inoculated at the highest dose was used for PrP and GFAP immunostaining, and hematoxylin and eosin (H&E) staining. Consistent with the features observed with typical 263K clinical disease, PrP deposition, astrogliosis (GFAP), and spongiform change were observed. Scale bar = 50 µm.

**Extended Data Fig. 2.**
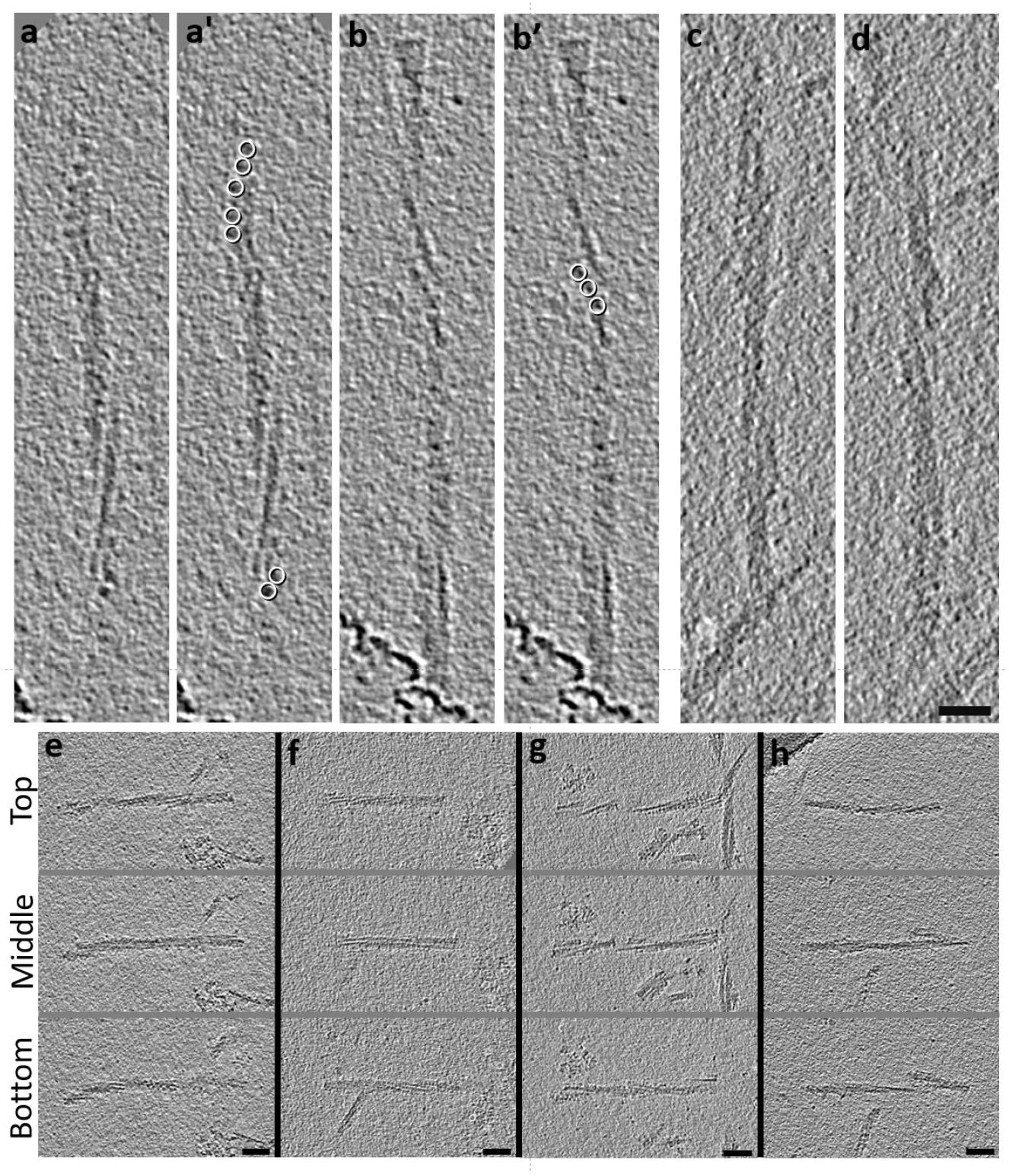
Tomography of 263K prions shows left‐handed fibrils and asymmetric decoration with globules. **a**, Tomographic slice of a fibril with globules visible on the ends, outlined with white circles in **a’. b**, Another slice with globules, as marked in **b’. c**,**d**, Two slices of a 263K fibril without apparent globules. **e‐h**, Fibrils from 4 different tomograms showing the top, middle, and bottom slices, illustrating the left‐handed helix found in >99% of fibrils analyzed by tomography. Scale bars: **a‐d** = 25 nm, **e‐h** = 50 nm.

**Extended Data Table 1.** Tomogram‐based dimensions of 263K fibrils of preps 1 & 2 with and without globules

| Fibril Prep | Globules | n | Length (nm) | Width (nm) |
| --- | --- | --- | --- | --- |
| 1 | + | 15 | 188 ±87 | 20 ±3 |
|  | - | 64 | 176 ±66 | 16 ±3 |
| 2 | + | 98 | 137 ±54 | 18 ±2 |
|  | - | 71 | 98 ±33 | 16 ±2 |

**Extended Data Table 2.** Cryo‐EM data, refinement, and validations

|  | 263K (pdb, emdb) | With 194-196 |
| --- | --- | --- |
| **Data collection and processing** |  |  |
| Magnification | 81,000x |  |
| Voltage (kV) | 300 |  |
| Electron dose (e-/ Å<sup>2</sup>) | 60 |  |
| Pixel size (super resolution) | 0.545 |  |
| Symmetry imposed | C1 |  |
| Initial particle segments | 337368 |  |
| Final particle segments | 15884 |  |
| Map resolution (Å) (FSC 0.143) | 3.14 |  |
| Helical rise (Å) | 4.874 |  |
| Helical twist (°) | -0.847 |  |
| Map sharpening *B* factor (Å<sup>2</sup>) | -30.98 |  |
| **Model Refinement** |  |  |
| Initial Model | *de novo* |  |
| R.M.S. deviations |  |  |
| Bond lengths (Å) | 0.011 | 0.011 |
| Bond angles (°) | 1.284 | 1.283 |
| MolProbity score | 1.26 | 1.35 |
| Clash score | 2.77 | 2.10 |
| Rotamer outliers (%) | 0 | 0 |
| Ramachandran plot |  |  |
| Favored (%) | 96.83 | 94.66 |
| Allowed (%) | 3.17 | 5.34 |
| Outliers (%) | 0 | 0 |
| EM Ringer score | 5.01 | 4.68 |
| Model vs. Data (CC) | 0.80 | 0.80 |

**Extended Data Fig. 3.**
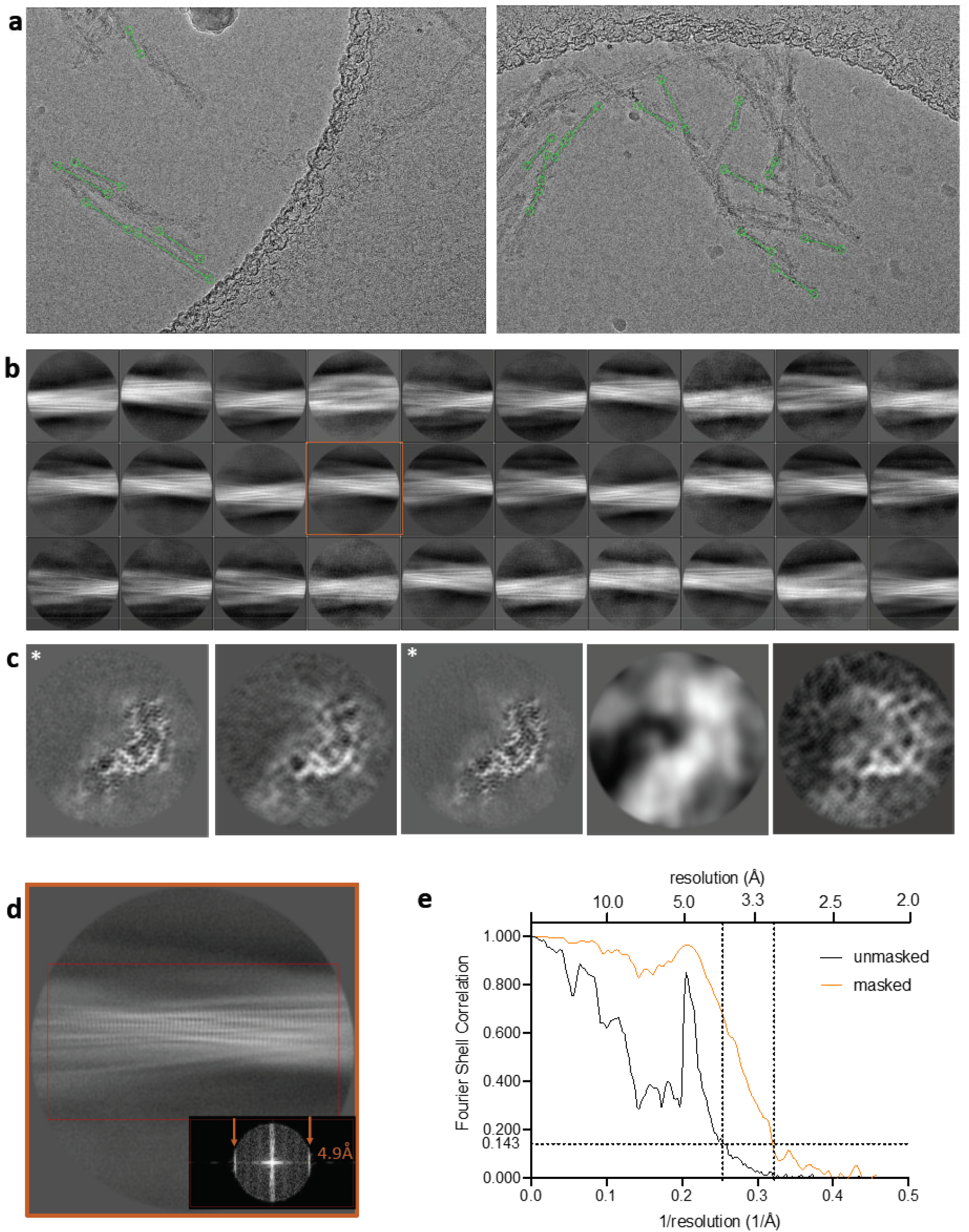
Particle selection and 2D and 3D classifications in Relion. **a**, Representative micrographs are shown with examples of manually selected segments of fibrils (green) used for particle extraction. **b**, Representative 2D classifications from a total of 35 used for 3D model building. **c**, Representative cross‐sections from 3D classes. The first and third classes (asterisks) were used for subsequent model refinement, and the others of this group discarded due to visually low resolution and poor alignment. **d**, Enlarged view of one of the 2D classes (highlighted in **b**) showing the 4.9Å repeated spacing perpendicular to the fibril axis. Associated fast Fourier transform indicating signals at 4.9 Å (inset). **e**, Fourier shell correlation plots of masked and unmasked models.

**Extended Data Fig. 4.**
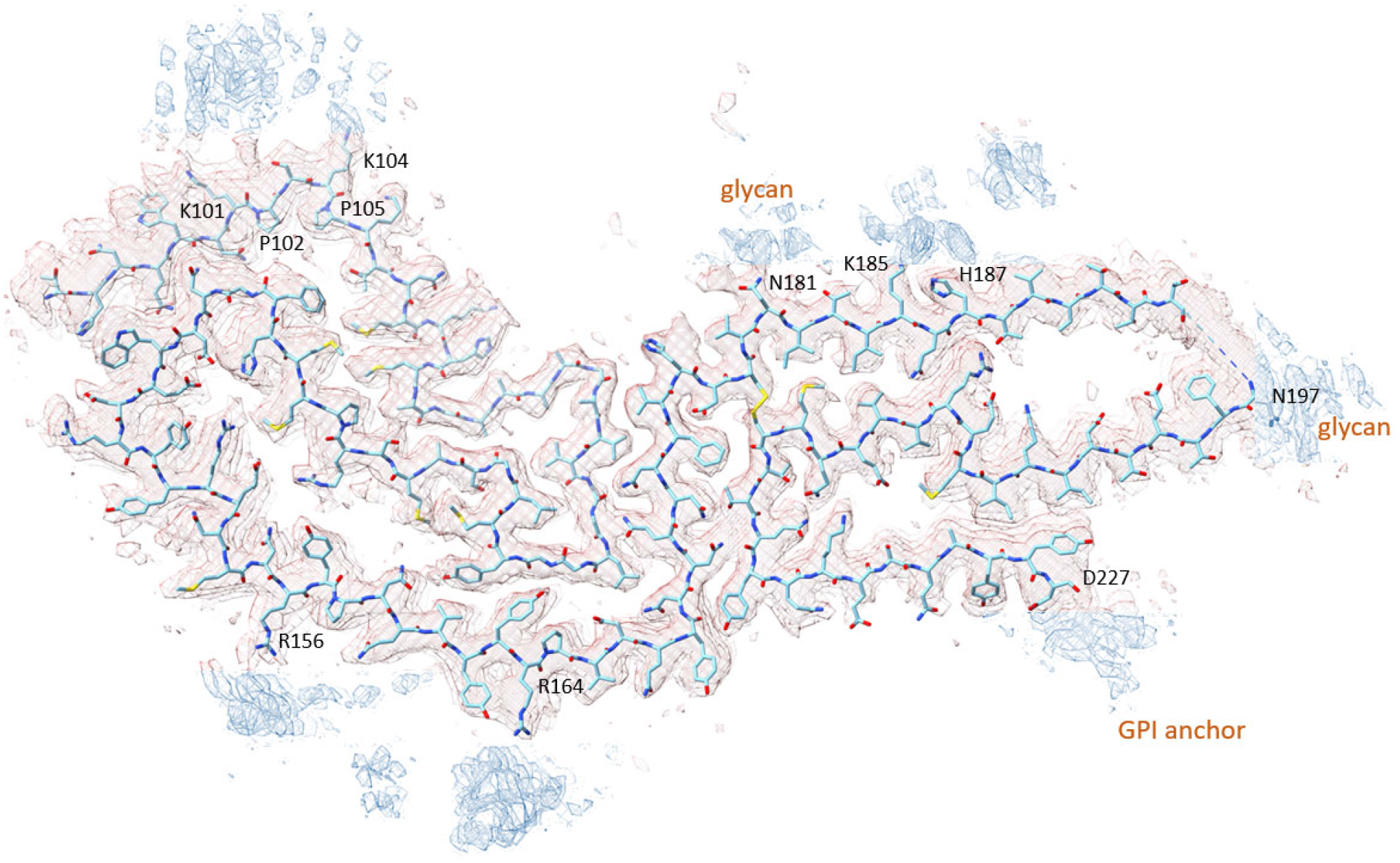
Peripheral map densities (blue) not occupied by the polypeptide backbone of residues 95‐227. Glycans and lipid anchor positions are labeled. Charged residues are often found next to unidentified peripheral map densities that are not associated with the known (labeled) glycan or GPI anchor attachment sites.

**Extended Data Fig. 5.**
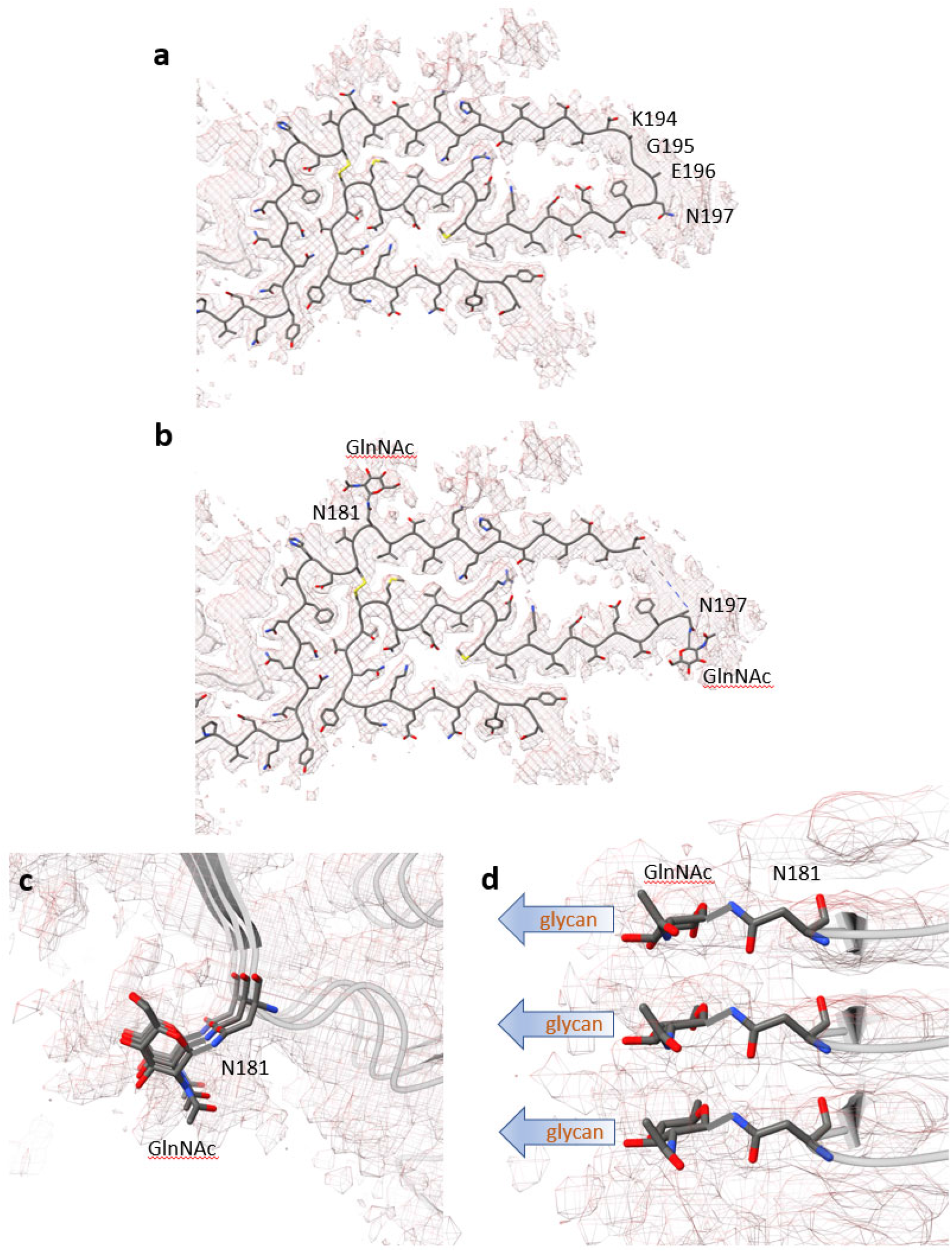
Additional models with residues K194‐E196 and first N‐acetylglucosamine on N181 and N197. **a**, A polypeptide backbone inclusive of residues K194‐E196 is shown within the map density (mesh). **b**, The first N‐acetylglucosamine (GlcNAc) units attached to the sidechains of N181 or N197 are shown within map densities. **c**, Magnified cross‐sectional view of N181 with the GlcNAc residue. **d**, Magnified side view of the model with GlcNAc residue placement in the map densities visible directly adjacent to N181. The remaining glycan extends off the fibril axis, and outside the resolved map densities.

**Extended Data Fig. 6:**
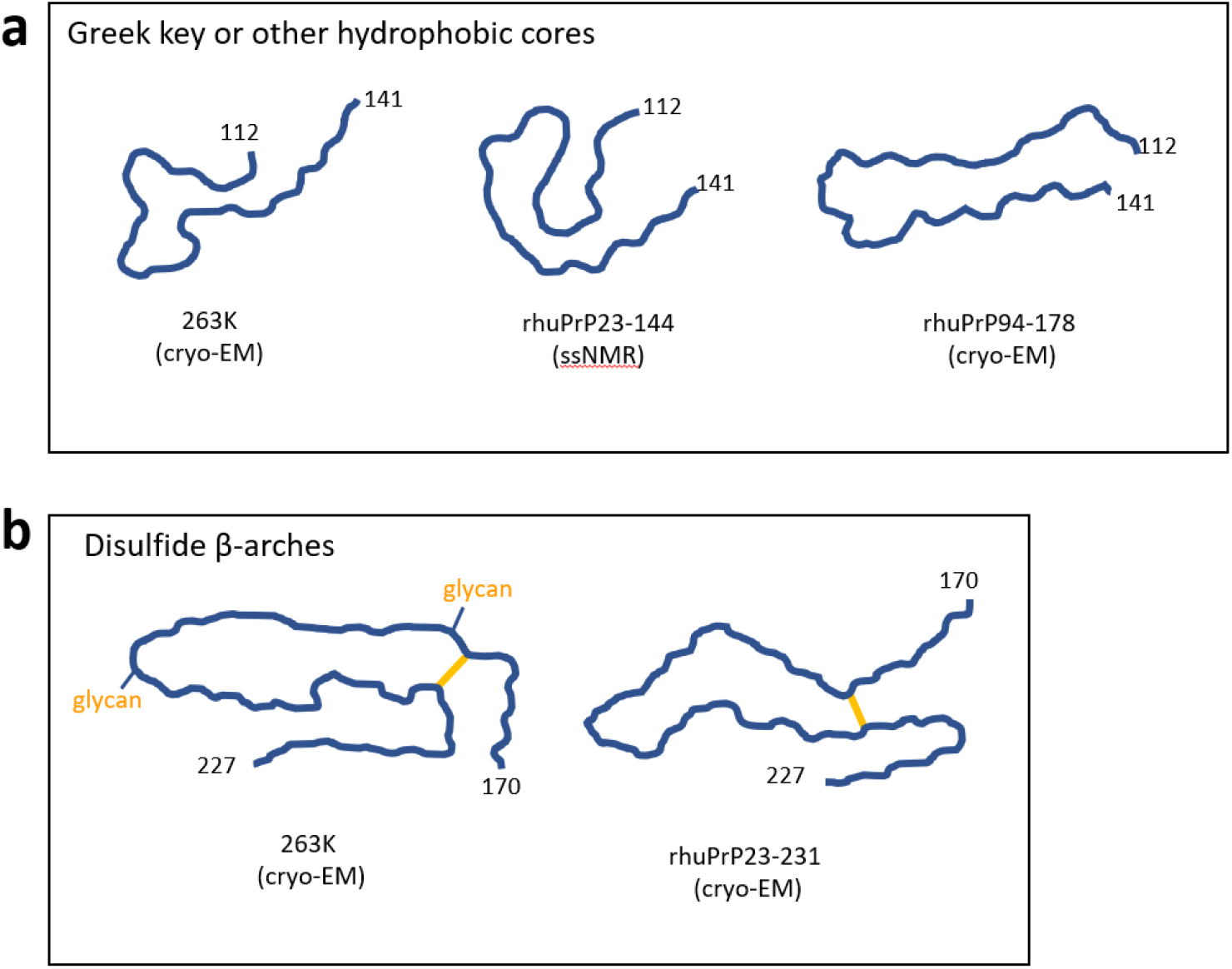
Comparison of 263K Greek key and disulfide β‐arches to related motifs in previously described recombinant PrP fibrils. **a**, Polypeptide backbone tracings of residues 112‐141 of the 263K prion Greek key compared to a Greek key‐like motif modelled previously for synthetic rhuPrP23‐144^46^ or a more extended hydrophobic core of rhuPrP94‐178^33^ fibrils based on solid state NMR or cryo‐EM data, respectively. **b**, Tracings of residues 170‐227 the cryo‐EM‐based disulfide β‐arches of 263K prions (current work) and synthetic rhuPrP23‐231 fibrils^34^. Yellow bar indicates the disulfide bond.

**Extended Data Fig. 7.**
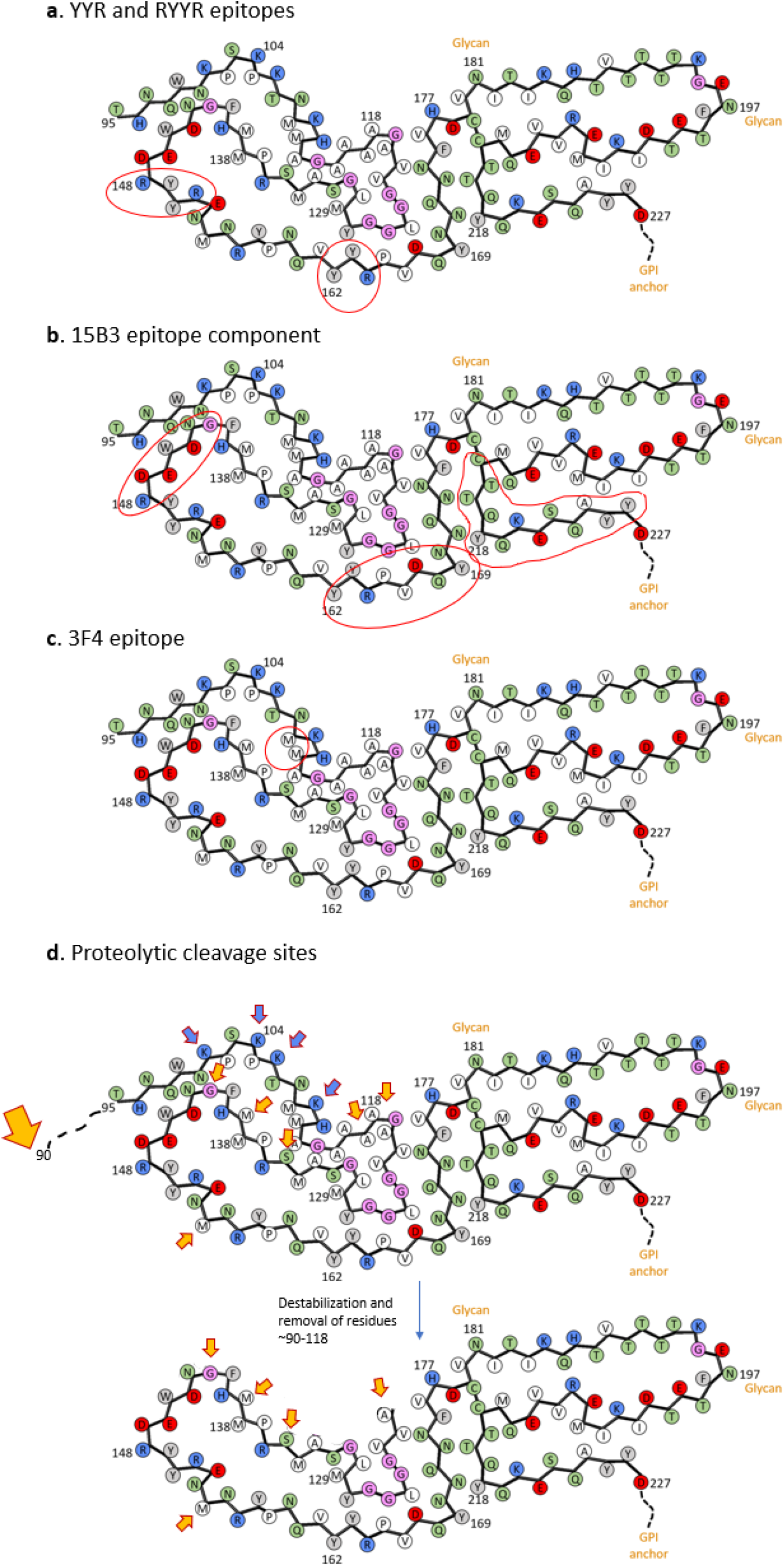
Antibody epitopes and proteolytic cleavage sites on 263K fibril core cross‐section. **a**, Exposure of YYR and RYYR epitopes (encircled in red), one or both of which are reactive with PrP^Sc^‐selective antibodies^66^. **b**, Locations of sequences that are thought to bracket elements of the PrP^Sc^‐ selective MAb 15B3 epitope (encircled in red). **c**, M109 and M112 (encircled in red), which are key elements of the MAb 3F4 antibody epitope. **d**, Primary (large yellow arrow) and conditional minor secondary (small yellow arrows) PK cleavage sites. Bottom depicts core that would remain after partial unfolding and removal of residues 90‐119, revealing the minor PK cleavage sites. Blue arrows mark lysines in this segment.

**Extended Data Fig. 8:**
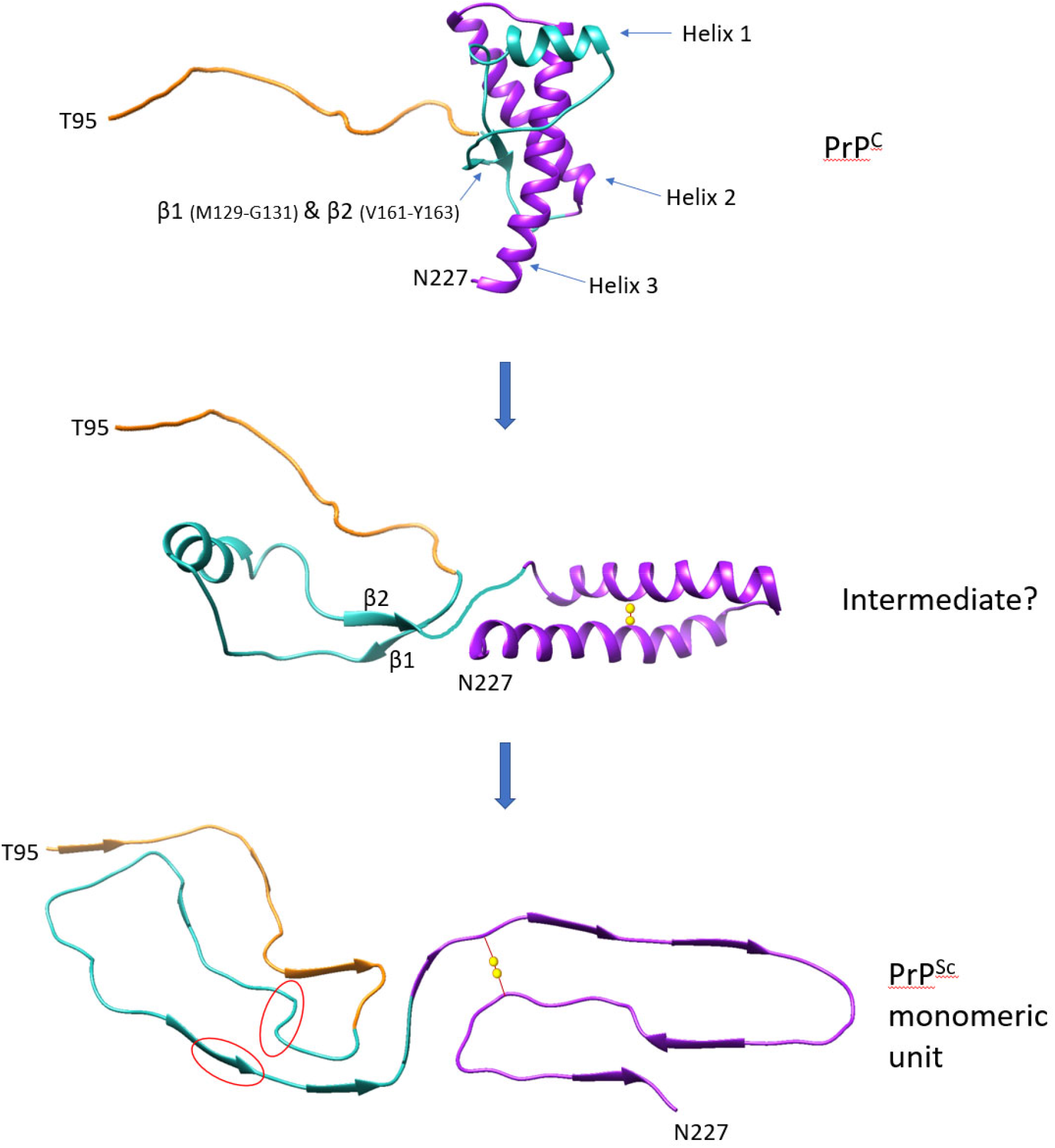
Domain separation and helix untwisting in conversion of PrP^C 67^ to PrP^Sc^. Models depicting the need for PrP^C^’s β1‐Helix 1‐β2 loop (turquoise) to separate from helices 2 and 3 (purple), and for each the helices to untwist into extended chains during conversion to PrP^Sc^. This mechanism is reminiscent of a previously proposed “banana peeling model”^68^. Also notable is the breakage of the intramolecular backbone H‐bonds of PrP^C^’s β1‐β2 sheet as they become intermolecular in PrP^Sc^ and no longer in the same sheet (new positions encircled in red). Residues 23‐124 of PrP^C^ are disordered in PrP^C 67^ and only residues 95‐124 are depicted (orange) in the top 2 panels. Although these conformational changes must occur during conversion, the order of these and other events, and whether an intermediate specifically like the one depicted ever exists at any individual step, are not known. The disulfide bond between C179‐C214 (yellow spheres) is visible only in the lower 2 panels. Note: elements of these models may not be accurately matched in scale.

## Main References

1. Caughey, B.W., et al. Secondary structure analysis of the scrapie-associated protein PrP 27-30 in water by infrared spectroscopy. Biochemistry 30, 7672–7680 (1991).

2. Caughey, B. & Kraus, A. Transmissibility versus Pathogenicity of Self-Propagating Protein Aggregates. Viruses 11(2019).

3. Stahl, N., Borchelt, D.R., Hsiao, K. & Prusiner, S.B. Scrapie prion protein contains a phosphatidylinositol glycolipid. Cell 51, 229–240 (1987).

4. Bolton, D.C., Meyer, R.K. & Prusiner, S.B. Scrapie PrP 27-30 is a sialoglycoprotein. J.Virol. 53, 596–606 (1985).

5. Manuelidis, L., Valley, S. & Manuelidis, E.E. Specific proteins associated with Creutzfeldt-Jakob disease and scrapie share antigenic and carbohydrate determinants. Proc.Natl.Acad.Sci.USA 82, 4263–4267 (1985).

6. Rudd, P.M., et al. Glycosylation differences between the normal and pathogenic prion protein isoforms. Proc.Natl.Acad.Sci.U.S.A. 96, 13044–13049 (1999).

7. Baskakov, I.V., et al. The prion 2018 round tables (I): the structure of PrP(Sc). Prion 13, 46–52 (2019).

8. Prusiner, S.B. Prions. Proc. Natl. Acad. Sci. U.S.A. 95, 13363–13383 (1998).

9. Kraus, A., Groveman, B.R. & Caughey, B. Prions and the potential transmissibility of protein misfolding diseases. Annu. Rev. Microbiol. 67, 543–564 (2013).

10. Safar, J., Roller, P.P., Gajdusek, D.C. & Gibbs, C.J. Jr., Conformational transitions, dissociation, and unfolding of scrapie amyloid (prion) protein. J. Biol. Chem. 268, 20276–20284 (1993).

11. Pan, K.-M., et al. Conversion of alpha-helices into beta-sheets features in the formation of the scrapie prion protein. Proc. Natl. Acad. Sci. USA 90, 10962–10966 (1993).

12. Kocisko, D.A., et al. Cell-free formation of protease-resistant prion protein. Nature 370, 471–474 (1994).

13. Bessen, R.A., et al. Nongenetic propagation of strain-specific phenotypes of scrapie prion protein. Nature 375, 698–700 (1995).

14. Telling, G.C., et al. Evidence for the conformation of the pathologic isoform of the prion protein enciphering and propagating prion diversity. Science 274, 2079–2082 (1996).

15. Caughey, B., Raymond, G.J., Kocisko, D.A. & Lansbury, P.T. Jr., Scrapie infectivity correlates with converting activity, protease resistance, and aggregation of scrapie-associated prion protein in guanidine denaturation studies. J. Virol. 71, 4107–4110 (1997).

16. Silveira, J.R., et al. The most infectious prion protein particles. Nature 437, 257–261 (2005).

17. Caughey, B., Baron, G.S., Chesebro, B. & Jeffrey, M. Getting a grip on prions: oligomers, amyloids, anchors and pathological membrane interactions. Annu. Rev. Biochem. 78, 177–204 (2009).

18. Wuthrich, K. & Riek, R. Three - dimensional structures of prion proteins. Adv.Prot.Chem. 57, 55–82 (2001).

19. Merz, P.A., Somerville, R.A., Wisniewski, H.M. & Iqbal, K. Abnormal fibrils from scrapie-infected brain. Acta Neuropathol. 54, 63–74 (1981).

20. Diringer, H., et al. Scrapie infectivity, fibrils and low molecular weight protein. Nature 306, 476–478 (1983).

21. Wille, H., et al. Structural studies of the scrapie prion protein by electron crystallography. Proc.Natl.Acad.Sci.U.S.A 99, 3563–3568 (2002).

22. Caughey, B., Kocisko, D.A., Raymond, G.J. & Lansbury, P.T. Aggregates of scrapie associated prion protein induce the cell-free conversion of protease-sensitive prion protein to the protease-resistant state. Chem.& Biol. 2, 807–817 (1995).

23. Sim, V.L. & Caughey, B. Ultrastructures and strain comparison of under-glycosylated scrapie prion fibrils. Neurobiol.Aging 30, 2031–2042 (2009).

24. Amenitsch, H., Benetti, F., Ramos, A., Legname, G. & Requena, J.R. SAXS structural study of PrP(Sc) reveals ∼11 nm diameter of basic double intertwined fibers. Prion 7, 496–500 (2013).

25. Vazquez-Fernandez, E., et al. The Structural Architecture of an Infectious Mammalian Prion Using Electron Cryomicroscopy. PLoS Path. 12, e1005835 (2016).

26. Spagnolli, G., et al. Full atomistic model of prion structure and conversion. PLoS Path. 15, e1007864 (2019).

27. Terry, C., et al. Structural features distinguishing infectious ex vivo mammalian prions from non-infectious fibrillar assemblies generated in vitro. Sci Rep 9, 376 (2019).

28. Cobb, N.J., Sonnichsen, F.D., McHaourab, H. & Surewicz, W.K. Molecular architecture of human prion protein amyloid: a parallel, in-register beta-structure. Proc Natl Acad Sci U.S.A 104, 18946–18951 (2007).

29. Cobb, N.J., Apetri, A.C. & Surewicz, W.K. Prion protein amyloid formation under native-like conditions involves refolding of the C-terminal alpha-helical domain. J. Biol. Chem. 283, 34704–34711 (2008).

30. Tycko, R., Savtchenko, R., Ostapchenko, V.G., Makarava, N. & Baskakov, I.V. The alpha-helical C-terminal domain of full-length recombinant PrP converts to an in-register parallel beta-sheet structure in PrP fibrils: evidence from solid state nuclear magnetic resonance. Biochemistry 49, 9488–9497 (2010).

31. Groveman, B.R., et al. Parallel in-register intermolecular beta-sheet architectures for prion-seeded prion protein (PrP) amyloids. J. Biol. Chem. 289, 24129–24142 (2014).

32. Theint, T., et al. Species-dependent structural polymorphism of Y145Stop prion protein amyloid revealed by solid-state NMR spectroscopy. Nat Commun 8, 753 (2017).

33. Glynn, C., et al. Cryo-EM structure of a human prion fibril with a hydrophobic, protease-resistant core. Nat. Struct. Mol. Biol. 27, 417–423 (2020).

34. Wang, L.Q., et al. Cryo-EM structure of an amyloid fibril formed by full-length human prion protein. Nat. Struct. Mol. Biol. 27, 598–602 (2020).

35. Kim, J.I., et al. Mammalian prions generated from bacterially expressed prion protein in the absence of any mammalian cofactors. J. Biol. Chem. 285, 14083–14087 (2010).

36. Choi, J.K., et al. Amyloid fibrils from the N-terminal prion protein fragment are infectious. Proc Natl Acad Sci U S A 113, 13851–13856 (2016).

37. Goedert, M., et al. Tau Protein and Frontotemporal Dementias. Adv. Exp. Med. Biol. 1281, 177–199 (2021).

38. Kraus, A., et al. PrP P102L and nearby lysine mutations promote spontaneous in vitro formation of transmissible prions. J. Virol. (2017).

39. Groveman, B.R., et al. Role of the central lysine cluster and scrapie templating in the transmissibility of synthetic prion protein aggregates. PLoS Path. 13, e1006623 (2017).

40. Prusiner, S.B., Groth, D.F., Bolton, D.C., Kent, S.B. & Hood, L.E. Purification and structural studies of a major scrapie prion protein. Cell 38, 127–134 (1984).

41. Hope, J., et al. The major polypeptide of scrapie-associated fibrils (SAF) has the same size, charge distribution and N-terminal protein sequence as predicted for the normal brain protein (PrP). EMBO J. 5, 2591–2597 (1986).

42. Wilham, J.M., et al. Rapid End-Point Quantitation of Prion Seeding Activity with Sensitivity Comparable to Bioassays. PLoS Path. 6, e1001217 (2010).

43. Artikis, E., Roy, A., Verli, H., Cordeiro, Y. & Caughey, B. Accommodation of In-Register N-Linked Glycans on Prion Protein Amyloid Cores. ACS Chem Neurosci 11, 4092–4097 (2020).

44. Spagnolli, G., et al. Modeling PrP(Sc) Generation Through Deformed Templating. Front Bioeng Biotechnol 8, 590501 (2020).

45. Tuttle, M.D., et al. Solid-state NMR structure of a pathogenic fibril of full-length human alpha-synuclein. Nat. Struct. Mol. Biol. 23, 409–415 (2016).

46. Theint, T., et al. Structural Studies of Amyloid Fibrils by Paramagnetic Solid-State Nuclear Magnetic Resonance Spectroscopy. J. Am. Chem. Soc. 140, 13161–13166 (2018).

47. Hafner-Bratkovic, I., et al. Globular domain of the prion protein needs to be unlocked by domain swapping to support prion protein conversion. J.Biol.Chem. 286, 12149–12156 (2011).

48. Jeffrey, M. Review: membrane-associated misfolded protein propagation in natural transmissible spongiform encephalopathies (TSEs), synthetic prion diseases and Alzheimer’s disease. Neuropathol. Appl. Neurobiol. 39, 196–216 (2013).

49. Tycko, R. & Wickner, R.B. Molecular structures of amyloid and prion fibrils: consensus versus controversy. Acc Chem Res 46, 1487–1496 (2013).

50. Donne, D.G., et al. Structure of the recombinant full-length hamster prion protein PrP(29-231): the N terminus is highly flexible [see comments]. Proc.Natl.Acad.Sci.U.S.A. 94, 13452–13457 (1997).

## Additional References for Methods, Extended Data and Supplemental Information

51. Race, R., Oldstone, M. & Chesebro, B. Entry versus blockade of brain infection following oral or intraperitoneal scrapie administration: Role of prion protein expression in peripheral nerves and spleen. J. Virol. 74, 828–833 (2000).

52. Raymond, G.J. & Chabry, J. Purification of the pathological isoform of prion protein (PrPSc or PrPres) from transmissible spongiform encephalopathy-affected brain tissue. in Techniques in Prion Research (eds. Lehmann, S. & Grassi, J.) 16–26 (Birkhauser Verlag, Basel, 2004).

53. Caughey, B., Raymond, G.J., Ernst, D. & Race, R.E. N-terminal truncation of the scrapie-associated form of PrP by lysosomal protease(s): implications regarding the site of conversion of PrP to the protease-resistant state. J.Virol. 65, 6597–6603 (1991).

54. Baron, G.S., et al. Effect of glycans and the glycophosphatidylinositol anchor on strain dependent conformations of scrapie prion protein: improved purifications and infrared spectra. Biochemistry 50, 4479–4490 (2011).

55. Mastronarde, D.N. Automated electron microscope tomography using robust prediction of specimen movements. J Struct Biol 152, 36–51 (2005).

56. Kremer, J.R., Mastronarde, D.N. & McIntosh, J.R. Computer visualization of three-dimensional image data using IMOD. J Struct Biol 116, 71–76 (1996).

57. Scheres, S.H.W. Amyloid structure determination in RELION-3.1. Acta Crystallogr D Struct Biol 76, 94–101 (2020).

58. Rohou, A. & Grigorieff, N. CTFFIND4: Fast and accurate defocus estimation from electron micrographs. J Struct Biol 192, 216–221 (2015).

59. Emsley, P., Lohkamp, B., Scott, W.G. & Cowtan, K. Features and development of Coot. Acta Crystallogr D Biol Crystallogr 66, 486–501 (2010).

60. Afonine, P.V., et al. Towards automated crystallographic structure refinement with phenix.refine. Acta Crystallogr D Biol Crystallogr 68, 352–367 (2012).

61. Headd, J.J., et al. Use of knowledge-based restraints in phenix.refine to improve macromolecular refinement at low resolution. Acta Crystallogr D Biol Crystallogr 68, 381–390 (2012).

62. Murshudov, G.N., Vagin, A.A. & Dodson, E.J. Refinement of macromolecular structures by the maximum-likelihood method. Acta Crystallogr D Biol Crystallogr 53, 240–255 (1997).

63. Williams, C.J. Duke University (2015).

64. Chen, V.B., et al. MolProbity: all-atom structure validation for macromolecular crystallography. Acta Crystallogr D Biol Crystallogr 66, 12–21 (2010).

65. Barad, B.A., et al. EMRinger: side chain-directed model and map validation for 3D cryo-electron microscopy. Nat. Methods 12, 943–946 (2015).

66. Paramithiotis, E., et al. A prion protein epitope selective for the pathologically misfolded conformation. Nat.Med. 9, 893–899 (2003).

67. James, T.L., et al. Solution structure of a 142-residue recombinant prion protein corresponding to the infectious fragment of the scrapie isoform. Proc.Natl.Acad.Sci.U.S.A. 94, 10086–10091 (1997).

68. Adrover, M., et al. Prion Fibrillization Is Mediated by a Native Structural Element That Comprises Helices H2 and H3. J. Biol. Chem. 285, 21004–21012 (2010).

69. Caughey, B., Raymond, G.J. & Bessen, R.A. Strain-dependent differences in beta-sheet conformations of abnormal prion protein. J.Biol.Chem. 273, 32230–32235 (1998).

70. Smirnovas, V., et al. Structural organization of brain-derived mammalian prions examined by hydrogen-deuterium exchange. Nat.Struct.Mol.Biol. 18, 504–506 (2011).

71. Speare, J.O., Rush, T.S., Bloom, M.E. & Caughey, B. The role of helix 1 aspartates and salt bridges in the stability and conversion of prion protein. J.Biol.Chem. 278, 12522–12529 (2003).

72. Korth, C., et al. Prion (PrPSc)-specific epitope defined by a monoclonal antibody. Nature 390, 74–77 (1997).

73. Orru, C.D., et al. Prion disease blood test using immunoprecipitation and improved quaking-induced conversion. mBio 2, e00078–00011 (2011).

74. Safar, J., et al. Eight prion strains have PrP(Sc) molecules with different conformations [see comments]. Nat.Med. 4, 1157–1165 (1998).

75. Sajnani, G., Pastrana, M.A., Dynin, I., Onisko, B. & Requena, J.R. Scrapie prion protein structural constraints obtained by limited proteolysis and mass spectrometry. J.Mol.Biol. 382, 88–98 (2008).

76. Groveman, B.R., et al. Charge neutralization of the central lysine cluster in prion protein (PrP) promotes PrP(Sc)-like folding of recombinant PrP amyloids. J. Biol. Chem. 290, 1119–1128 (2015).

77. Wong, C., et al. Sulfated glycans and elevated temperature stimulate PrP^Sc^ dependent cell-free formation of protease-resistant prion protein. EMBO J. 20, 377–386 (2001).

78. Kocisko, D.A., Lansbury, P.T. Jr., & Caughey, B. Partial unfolding and refolding of scrapie-associated prion protein: evidence for a critical 16-kDa C-terminal domain. Biochemistry 35, 13434–13442 (1996).

79. McKenzie, D., et al. Reversibility of scrapie inactivation is enhanced by copper. J.Biol.Chem. 273, 25545–25547 (1998).

80. Eisenberg, D.S. & Sawaya, M.R. Structural Studies of Amyloid Proteins at the Molecular Level. Annu. Rev. Biochem. 86, 69–95 (2017).

