## Supplementary Information for "Structure of an infectious mammalian prion"

### Supplementary Movies

**Movie 1: Globules_SB25nm.mp4**
Movie through a tomogram of a 263K fibril that is decorated with globules along one side of the fibril. Globules are encircled in white. The tomogram was filtered using nonlinear anisotropic diffusion to aid in visualization. Z-slices are ~1 nm. Scale bar = 25 nm.

**Movie 2: NoGlobules_SB25nm.mp4**
Movie through a tomogram of a 263K fibril that lacks apparent globules. The tomogram was filtered using nonlinear anisotropic diffusion to aid in visualization. Z-slices are ~1 nm. Scale bar = 25 nm.

**Movie 3: WholeTomo_SB100nm.mp4**
Movie through a representative tomogram showing 263K fibrils in different orientations within the ice. All fibrils are left-handed and some are decorated with globules. The tomogram was filtered using nonlinear anisotropic diffusion to aid in visualization. Z-slices are ~1 nm. Scale bar = 100 nm.

### Supplementary Discussion

#### Does the final model represent the $PrP^{Sc}$ preparation as a whole?

A number of lines of evidence suggest that our final high-resolution structure represents the bulk of our highly infectious $PrP^{Sc}$ fibril preparations: 1) Negative stain transmission EM and low magnification imaging of the entire $PrP^{Sc}$ preparation showed consistent fibril morphologies across many grids with little non-fibrillar material; 2) Intermediate resolution (~8 Å) structures inclusive of >240,000 particles were largely comparable in overall morphology to the 3.1 Å map achieved by disposing of particles that were of insufficient quality due to reasons such as poor ice thickness, oblique (z-axis) fibril image angles, or fibril overlaps that might have been unrecognized in the manual fibril selection process; 3) This selection process involved picking any visible and apparently individual fibril segment, and was not otherwise selective for any particular morphology; 4) 2D and 3D classifications only indicated a single fibril morphology even prior to extensive distillation of particles to curate those of sufficient quality to give 3.1 Å resolution; 5) Biochemical analyses of the $PrP^{Sc}$ preparations were consistent with extensive published data indicating one predominant PK-resistant 263K core comprising residues ~90-231[40,41] which, in turn, is compatible with the extent of the highly resolved amyloid core of our model.

**Extended comparison of 263K prion model to published data**

*Secondary structure composition*

Early Fourier transform infrared spectroscopy measurements of PK-treated 263K, or closely related, hamster prion scrapie $PrP^{Sc}$ indicated that it was roughly half β-sheet, and 21-31% turns or random coils[1,11,69]. Although the remainder (17-21%) was interpreted to be α-helix in these initial infrared studies, early circular dichroism[10] and more recent infrared[54,70] spectroscopy studies have indicated that PK-treated $PrP^{Sc}$ preparations lack α-helical content altogether. Our current findings confirm the latter conclusion for 263K $PrP^{Sc}$ fibrils. The ability of helices 2 and 3 to convert to β-sheet in the formation of amyloid fibrils has been demonstrated previously[29-31,34,68]. To our knowledge, conversion to β-sheet has not been shown previously for helix 1 but destabilization of this helix by elimination of its internal salt bridges promotes $PrP^{C}$ conversion to $PrP^{Sc}$ [71].

*Exposure of antibody epitopes*

One orthogonal means of testing our model is to ask if the epitopes of antibodies that are known to bind, or not bind, to $PrP^{Sc}$ are accessible on the fibril surface. Antibodies that specifically bind to $PrP^{Sc}$, and not $PrP^{C}$, are rare, but prominent examples are YYR (or RYYR) antibodies[66] and the monoclonal IgM 15B3[72] (Extended Data Fig. 7a). YYR epitopes are represented twice within residues 148-151 and 162-164[66]. Although the 162-164 epitope appears to be blocked by an unidentified density, the 148-151 (RYYR) epitope appears to be exposed. In both cases the sidechain of the middle Y residue is not exposed, but the previous YYR antibody study concluded that "access to the terminal tyrosine and arginine residues may be more important than the central tyrosine"[66].

Elements of the discontinuous conformational MAb 15B3 epitope(s) are contained within residues 142-148, 162-170, and 214-226[72]. At least substantial portions of these sequence ranges (namely, residues 146-148, 162-170 and 218-226) are presented on conformationally contiguous strands on the same side of the 263K fibril (Extended Data Fig. 7b). Although these epitope components are not immediately contiguous in the 263K structure, we note that MAb 15B3 is an IgM with 10 antigen combining sites per molecule. This, together with the repetition of the antigenic sites along the fibril, might allow high avidity binding that is based on a multiplicity of relatively low affinity interactions with individual components of the apparently conformational epitope. The fact that these antibodies are known to bind to multiple strains of $PrP^{Sc}$, including those of humans[72,73], is consistent with the possibility that this relative alignment of these epitope sequences on the $PrP^{Sc}$ surface is not restricted to 263K prions.

Another monoclonal antibody, 3F4, binds to $PrP^{C}$, but is buried in $PrP^{Sc}$ unless it is at least partially denatured[74]. Consistent with the latter observation, M109 and 112, which are key component of the 3F4 epitope, are buried within the 263K structure (Extended Data Fig. 7c).

*Known PK cleavage sites*

Although intact 263K prion fibrils are readily cleaved by PK before G90 (Extended Data Fig. 7d; big yellow arrow), other much more minor proteolytic (PK or trypsin) cleavage sites (small yellow arrows) have been observed in the regions of residues 117-119, 135, 139 and 142 and 154[75]. Residues 117-119 and 154 are exposed to solvent on the fibril surface. Although residues 117-119 lie at the base of a shallow groove that would likely be partially obscured by glycans on N181 (Fig. 4), they might become more exposed if flexing about a potential hinge at residues 165-170 allowed the N-terminal domain of residues ~90-168 to peel away from the remaining C-terminal domain. Residues 135, 139 and 142 line a pocket that would be exposed by cleavage at 117-119 and removal of a sub-domain formed by residues ~97-116. Notably, this sub-domain contains the cationic lysine cluster (blue arrows) that is suspected of binding polyanionic ligands such as sulfated glycosaminoglycans that are structurally heterogeneous[38,39,76]. Such non-protein ligands could protect this domain from proteolysis, but occasional gaps in this protection might allow cleavages in this sub-domain and exposure of these minor additional cleavage sites within residues 131-142. Importantly, it has long been known that the sulfated glycosaminoglycan, heparan sulfate, can promote the conversion of $PrP^{C}$ to a PK-resistant state[77], and that 2.5-3 M guanidine-HCl treatments can partially unfold 263K $PrP^{Sc}$ and/or dissociate protective cofactors. These effects might allow PK cleavage in this region[78,79].

*Lack of regular 10-Å inter-sheet spacings*

Consistent with previous studies of prion fibrils[25], we did not see a strong ~10 Å signal on the equator of either the Fourier transforms of single particle micrographs or representations from 2D classifications. Typically, in x-ray diffraction and cryo-EM analyses of amyloid fibrils, evidence of 10 Å spacings in the cross-section is seen that are due to common 10 Å face-to-face distances between opposed β-sheets[80]. However, in the 263K prion structure the spacings between β-sheets in the cross-section tend to be irregular, and often much wider, than is typical of other amyloid fibrils (Figs. 1&2). This variable spacing may account for the lack of a clear equatorial 10 Å signal in Fourier transformed power spectra from 263K cryo-EM images.

*H/D exchange data*

Mass spectrometry-based analyses of the rate of exchange of amide protons (H) of the PrP backbone with deuterons (D) from the solvent have shown that GPI-anchorless, N-linked glycan-deficient murine prion strains have greatly restricted exchange relative to $PrP^{C}$ throughout their PK-resistant cores[70]. Although comparable analyses of 263K prions have not been reported, and would be severely undermined by the presence of glycans and GPI anchors[70], the fact the many amide protons within the amyloid core are engaged in intermolecular hydrogen bonds with adjacent monomers would be consistent with slow rates of backbone H/D exchange seen in other fully infectious prions.

**REFERENCE LIST IS PROVIDED WITH MAIN MANUSCRIPT**
